# Single-cell proteomic maps of human induced pluripotent stem cells and their differentiation into motor neurons

**DOI:** 10.64898/2026.05.29.728893

**Authors:** Vladimir A. Zhemkov, Aleksandra Binek, Ali Haghani, Edo Israely, Shaughn Bell, Alba Sansa, George Lawless, Jennifer Van Eyk, Clive N. Svendsen

## Abstract

The majority of single cell-studies use RNA to identify the cell state. However, many RNAs are transient and decrease in response to elevated protein to control homeostasis. Thus, final cell states are largely defined by their unique and dynamic protein composition not RNA. Here a single-cell proteomics approach was used to identify the proteomic profile of human induced pluripotent stem cells (iPSCs) before and during their *in vitro* differentiation to motor neurons. By measuring up to 1000 proteins in each cell, novel clusters of growing iPSCs could be characterized along with a new proteomic pathway that defines motor neuron development. Interestingly, there was a dynamic and cell type-specific discordance between the protein levels and their corresponding messenger RNAs. This lays the foundation for drawing new single-cell proteomic maps of developing human tissues.

**IN BRIEF:** In this article, we report the first single-cell proteomic maps of induced pluripotent stem cells (iPSCs) and their differentiation into motor neurons (MNs). By identifying proteomes of individual cells, we resolve iPSC and MN states, their proteomic, metabolomic and organellar heterogeneity. We show considerable, stage-specific discordance between the transcriptome and the proteome in differentiated neurons.

## INTRODUCTION

Human induced pluripotent stem cells (iPSCs) can be differentiated into neuronal subtypes, providing a powerful *in vitro* model of neurodevelopment and neurodegenerative disorders^1,2^. In particular, spinal and cortical motor neurons (MNs) are critical to neurodegenerative disorders such as ALS and SMA as well as injuries^1,2^. Advances in single-cell/single-nuclei RNA sequencing (sc/sn RNA-seq) approaches highlight the previously under-appreciated heterogeneity of the RNA composition, or the transcriptome, of spinal cord MNs *in vitro* and *in vivo*^3–8^. However, RNA-based approaches miss the complexity of the MN proteomic composition and architecture.

Cellular function and homeostasis are dictated by the composition of protein networks, or the proteome. Importantly, large-scale datasets that profiled bulk RNA and bulk protein composition of iPSC-derived MNs^9–11^ highlighted significant discrepancies between the cellular transcriptome and proteome in brain tissue and cultured neurons. However, as bulk proteomic measurement attributes the total protein abundance to a heterogeneous cell population, it cannot resolve cell type-specific features or subpopulations within one cell type^10,12^. Further, unbiased determination of the MN proteome and their changes are particularly limited due to MN sparsity and signal dilution of the bulk methods^13–15^.

Shortcomings of bulk protein analysis can now be resolved with technological advancements to apply mass spectrometry (MS) at the single-cell proteomics (SCP) level^13,14,16–27^. Here we use our recently developed high-sensitivity MS-SCP based approach^16,17,28^ to characterize the proteomic heterogeneity of human iPSCs and iPSC-derived developing MNs.

## RESULTS

### Proteomic heterogeneity of induced pluripotent stem cell states

Many groups and consortium-led efforts have highlighted significant variations in iPSC responsiveness to developmental cues, lineage specifications and ultimately differentiation outcomes, which could in part relate to heterogeneity within the initial iPSC colonies^29–31^. While iPSCs often satisfy the general tests for pluripotency, the reasons for within and between-line heterogeneity remain puzzling and could have profound effects as science is shifting toward human-based disease models^32^.

By applying the first-generation MS-SCP approach, we recently reported proteomic heterogeneity of iPSC cultures^17^. Here, we again labeled cells with a viability dye and isolated single cells by fluorescence-activated cell sorting^16,17,28^ followed by analysis using our next generation MS-SCP instrument (**Fig 1 A**). We also included several pooled samples of 50 individual cells used for the data-independent acquisition-MS (DIA-MS) as the spectral library build^17^. To determine the proteomics of single cells, we used our “one pot” label-free single-cell proteomic approach combined with parallelized nanoflow dual-trap single-column liquid chromatography (nano-DTSC) (**Fig 1 A**)^16,33^. Peptide data at 1% FDR was merged, filtered, and normalized across wells. When all timepoints were combined, 2689 unique proteins were identified from a total 309 single iPSCs that underwent MS-SCP analysis with 1463 mean detected proteins per cell (**Fig. 1 B**).

**Figure 1.**
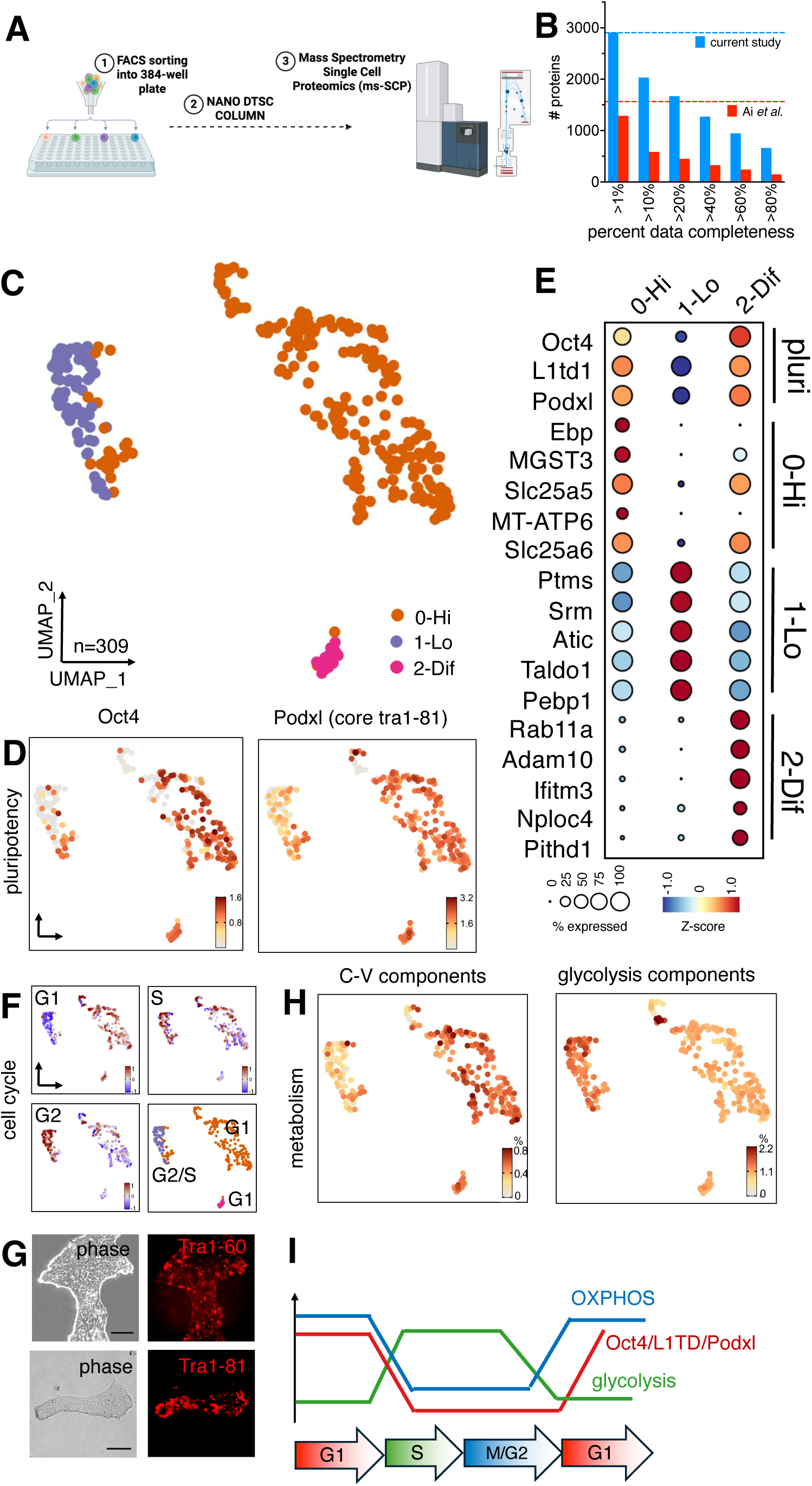
Mass-spectrometry single-cell proteomic profiling reveals heterogeneity of initial iPSC colonies. (**A**) The outline of the mass spectrometry – single-cell proteomics (MS-SCP) pipeline. iPSCs were dissociated into single-cell suspension and FACS-sorted into 384-well plates (1), including a 50-cell pools. Each cell was digested with trypsin, peptides were separated on nanoflow dual-trap single-column liquid chromatography (nano-DTSC) configuration (*16*) (2) and analyzed on a timsTOF Ultra 2 mass spectrometer at the single-cell level (3). (**B**) Data completeness of the generated iPSC dataset cross-compared to iPSC dataset from Ai *et al.* (*17*). (**C**) UMAP visualization of MS-SCP data for n=309 iPSCs reveals sub-clustering within the initial iPSC population. Three identified subclusters were labeled as labeled as 0-Hi (orange), 1-Lo (purple), 2-Dif (pink). (**D**) Abundance of pluripotency transcription factor Oct4 and surface iPSC marker podocalyxin (Podxl) are highly variable at the protein level in single cells. (**E**) Dot plot showing protein abundance and % of expressing cells for the canonical pluripotency markers and top five differentially abundant proteins enriched in each subcluster (increasing Z-score from blue to red; the size of bubble represents the percentage of expressing cells). (**F**) Assignment of cell cycle stage of the cells (increasing cell cycle modules from blue to red). (**G**) Heterogeneity of glycosylated Podxl antigens Tra1-60 and Tra1-81 by immunocytochemistry analysis of iPSC colonies. Scale bar = 100 mm. (**H**) Metabolic pathways were scored by calculating the sum of proteins within each MitoCarta pathway and normalized by total protein abundance to derive the percentage of each enzymatic pathway in each cell. Shown are percentages mitochondrial ATP synthase complex V (C-V) components and of the glycolytic pathway projected on the UMAP. (**I**) Summary of the pluripotency and metabolic fluctuations during cell cycle progression revealed by MS-SCP.

We first examined the proteomic landscape of single iPSCs. Compared with our previous approach^17^, the next generation instrument used in the current study provided a ∼20% increase in sensitivity^34^. This provided a significant increase (approximately 175%) in the detection of low abundant unique proteins (**Fig 1 B**). A more detailed cross-comparison of the current dataset to that reported by Ai *et al.*^28^ showed much higher numbers of proteins per cell (2689 verses 1534, respectively) (**Sup Fig 1 A**) permitting a more detailed description of individual iPSCs. Data integration with Harmony (*35*) using common highly variable proteins and accounting for batch and the number of detected proteins still resulted in distinct cellular populations (**Sup Fig 1 B**). For example, uniquely detected proteins in the current dataset included canonical stemness transcription factors such as Oct4 or alkaline phosphatase that were not detected by Ai *et al.*, in contrast to Lin28a (**Sup Fig 1 C**).

The protein abundance showed linearity between the two studies in the moderate-to-high intensity range, while in the low intensity range the current platform showed higher sensitivity (**Sup Fig 1 D**). Overall Spearman correlation indicated high consistency across different platforms with R=0.80.

Unbiased clustering of cells yielded three populations of iPSCs (**Fig 1 C)**. Interestingly, iPSCs expressed canonical pluripotency markers such as Oct4, Lin28A and L1TD1 at highly variable levels (**Fig 1 D and Sup Fig 2 A, B**). Specifically, cluster 0 (“0-Hi”) was characterized by high quantities of pluripotency markers Oct4, L1TD1 and podocalyxin (Podxl) - a known surface marker for iPSCs and embryonic stem cells (ESCs)^36^, while cluster 1 (“1-Lo”) had diminished expression of these markers (**Fig 1 D and Sup Fig 2 B**). Cluster 2 (“2-Dif”) expressed E-cadherin (E-cadh) and beta-catenin (b-Cat), consistent with early differentiation towards an epithelial state (**Sup Fig 2 C**). iPSCs also broadly expressed fibroblast growth factor 2, a paracrine signaling molecule involved in maintaining the pluripotent state (**Sup Fig 2 B**). Analysis of differentially abundant proteins highlighted distinct proteomic profiles between the “0-Hi” and “1-Lo” clusters as well as partially differentiated cells **(Fig. 1 E and Sup Table 1)**.

**Figure 2.**
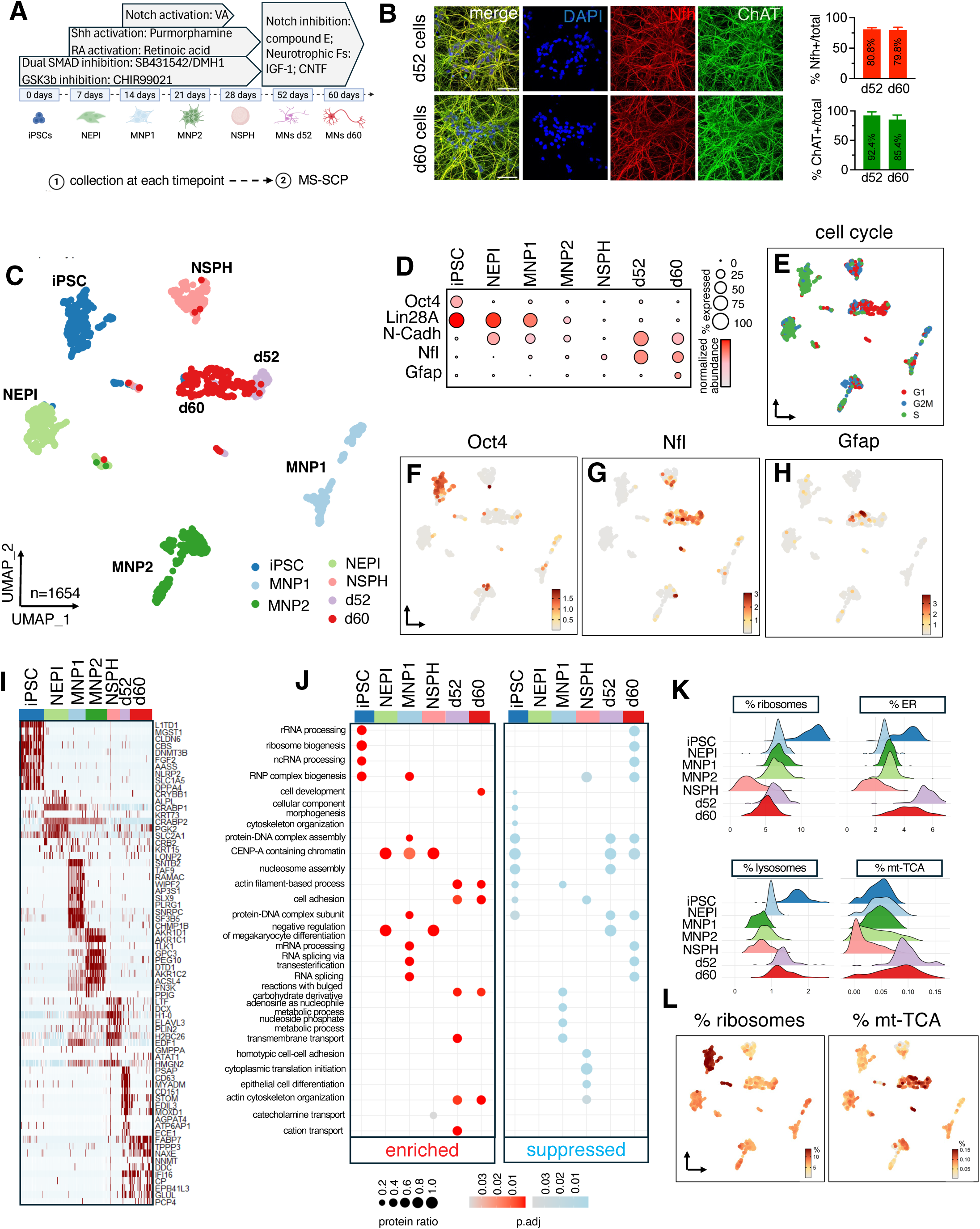
Mass-spectrometry single-cell proteomic profiling of differentiating motor neurons. (**A**) iPSCs were differentiated into neuroepithelial cells (NEPI), progenitors (MNP1/2 and NSPH) and MNs by exposure to small molecule morphogens. At each stage, cells were dissociated (1), processed and analyzed by MS-SCP as described above (2). (**B**) Morphology of differentiated cultures at d52 and d60 was assessed by immunocytochemistry analysis of neurofilament heavy (Nfh) (red), and choline acetyltransferase (ChAT) (green), with DAPI nuclear counterstain (blue). Quantification for the percent of each marker is shown per total cell. Scale bar = 50 mm. (**C**) UMAP visualization of MS-SCP data for n=1654 cells through the differentiation process from iPSC (blue) through NEPI (light green), MNP1 (light blue), MNP2 (green), NSPH (pink), d52 (purple) to MNs at d60 (red). (**D**) Dotplot showing the protein abundance and % of expressing cells for the pluripotency markers Oct4 and Lin28a; N-Cadherin (N-Cadh); MN-specific neurofilament light chain (Nfl) and glial-specific glial fibrillary acidic protein (GFAP) at each stage of differentiation (normalized abundance increasing from light to dark red; the size of bubble represents the percentage of expressing cells). (**E**) Assignment of cell cycle stage based on the levels of cycle-specific proteins. (**F-H**) Feature plots showing normalized abundance of Oct4, Nfl and GFAP at the single-cell level throughout the differentiation. (**I**) Heatmap plot showing normalized abundance for top 10 differentially enriched proteins at each differentiation stage (increasing Z-score from blue to red). (**J**) Gene set enrichment analysis (GSEA) of enriched (in red) and suppressed (in blue) biological processes associated with each stage (see associated legend for adjusted p-value scale bar and bubble size for protein ratios in each term). (**K**) Ridge plot showing cellular percentage of ribosomes, endoplasmic reticulum (ER), lysosomes and TCA enzymes (mt-TCA) at each stage. (**L**) Cellular percentage of ribosomes and TCA enzymes at the single-cell level, plotted as % of the proteins related to each pathway and shown on UMAP.

To further explore these three iPSC groups, we assessed cycle-dependent protein markers identified using single-cell proteomics analysis^37^, which led to classification into three states - G1, S, and G2 (**Fig 1 F**). The highest expression of Oct4 and Podxl was detected during the G1 cell cycle phase while their levels decreased significantly during the S and G2 phases (**Fig 1 D and 1 F**). Staining iPSC colonies for Tra1-60/Tra1-80, corresponding to glycosylated forms of Podxl, confirmed the presence of Podxl protein at heterogeneous levels consistent with the MS-SCP approach (**Fig 1 G)**. Given that the metabolic state of stem cells is important for maintaining pluripotency^38^, we next measured cumulative abundance of enzymes of the electron transport chain complexes, glycolysis and the TCA cycle. We observed distinct changes in cellular metabolism linked to cell cycle stage, with an increase in oxidative phosphorylation (OXPHOS, complex V-ATP synthase and the TCA cycle) in G1 and an increase in glycolysis in S/G2 phase (**Fig 1 H)**. Consistent with our single-cell data, a cycle-dependent shift in biosynthetic and mitochondrial proteins and cycle-dependent post-transcriptional regulation has been previously characterized in synchronized HeLa cells^39^ and multi-modal imaging approaches^40^.

Reports show that stem cells have shortened G1 phase^41^ and that the cell cycle stage can be important for pluripotency maintenance^42^ ^43^ and their sensitivity to developmental cues, ultimately influencing differentiation outcomes^43–45^. However, while several groups have performed RNA-seq analysis between multiple lines^32,46,47^, such clear within-colony heterogeneity of these pluripotency factors has not yet been detected using single-cell transcriptomics. At the protein level, several surface protein markers such as Tra1-60, SSEA3/4 and pluripotent transcription factors have been shown to be heterogeneous in ESCs and iPSCs using antibody staining and reporter lines^45,48–50^. Now the MS-SCP approach allowed us to better understand and characterize state transitions in mixed cell cultures and previously underappreciated differences in pluripotent markers related to cell stage and metabolism (summarized **Fig 1 I**). Overall, our comprehensive dataset suggests the role for more complex post-transcriptional regulation of the pluripotent factors during the S phase and the maintenance of pluripotency in general^51^ than previously observed.

### The proteomic landscape of differentiating human motor neurons

Having characterized the single-cell proteomic profile of iPSCs, we next interrogated proteomic changes as cells were differentiated towards a motor neuron lineage. The starting iPSCs were differentiated using our published 60-day protocol involving dual SMAD inhibition, caudalization and ventralization with developmental morphogens^52^ (**Fig. 2 A**). At each differentiation timepoint, cells were dissociated and analyzed by MS-SCP as described above. Besides the initial iPSC stage, cells were collected at the neuroepithelium (NEPI, day 7), early MN progenitor 1 (MNP1, d14), late MN progenitor 2 (MNP2, d21), and free floating neurosphere (NSPH, d28) stages (**Fig. 2 A**). For the final differentiation into MNs, neurospheres were plated and developed into more mature MNs and collected at days 52 and 60 (d52/d60) (**Fig. 2 A**). Differentiated d52/d60 cells had characteristic MN morphology and the vast majority expressed canonical MN markers, neurofilament heavy (Nfh) protein and choline acetyltransferase (ChAT) (**Fig 2 B**) and the transcription factor MNX1/Hb9 (**Sup Fig 3**). When all timepoints were combined, 3653 unique proteins were identified from a total 1634 single cells that underwent MS-SCP analysis with 930 mean detected proteins per cell.

**Figure 3.**
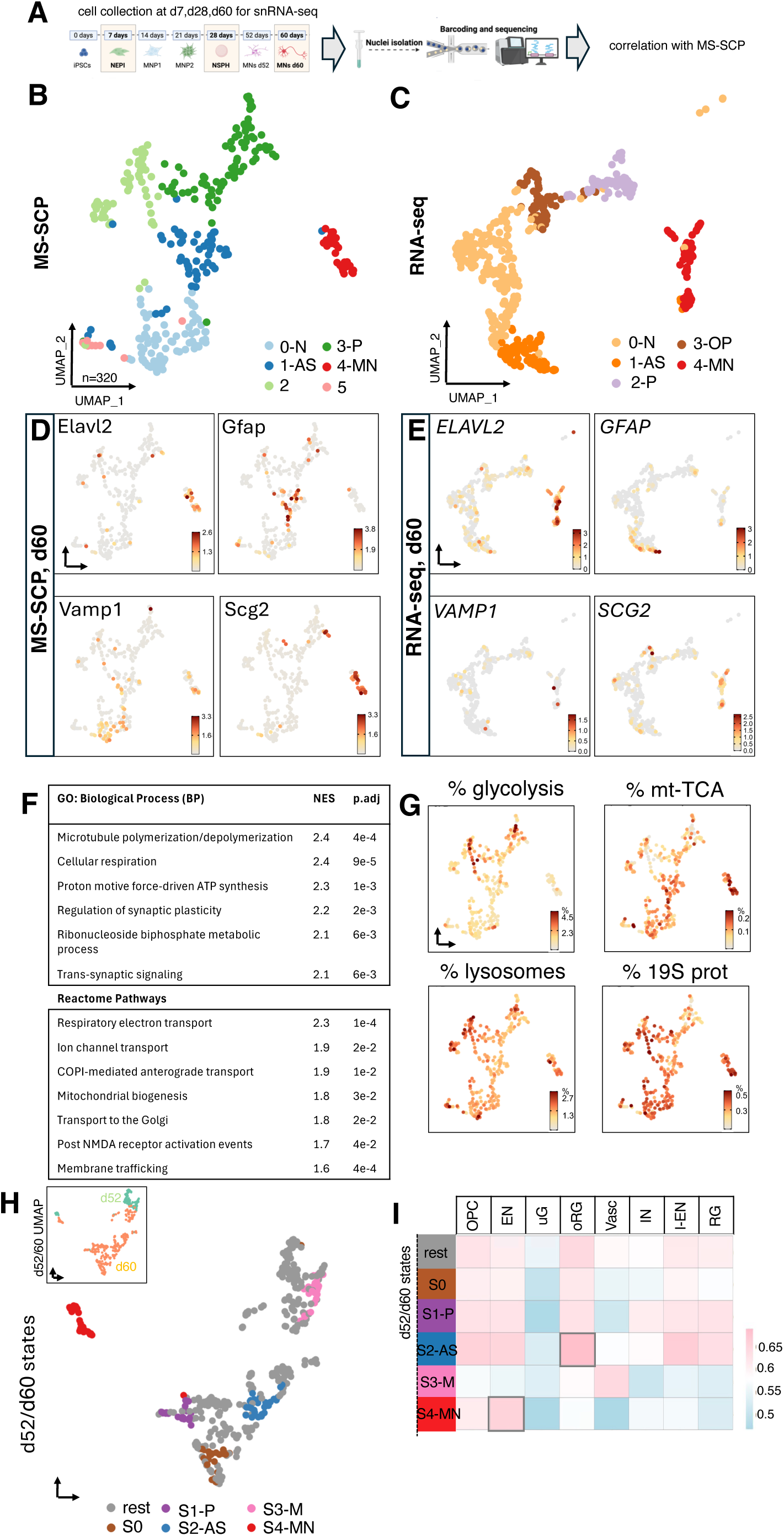
Identification of MN and glial clusters using MS-SCP of day 60 cultures. (**A**) To complement MS-SCP analysis, paired snRNA-seq of selected differentiation timepoints was performed to aid in cell type identification and investigation of the RNA-protein discordance. (**B**) Identification of several distinct cellular clusters of in vitro-derived d60 MN cultures using MS-SCP. Identified clusters include: “0-N” neuronal cluster (light blue), “1-AS” astrocyte cluster (blue), “2” (light green), “3-P” proliferative/neural stem cell cluster (dark green), “4-MN” motor neurons (red) and “5” (pink). (**C**) Identification of cellular clusters by snRNA-seq. Identified clusters include: “0-N” neuronal cells (light brown), “1-AS” astrocytes (orange), “2-P” proliferative/neural stem cells (violet), “3-OP” oligodendrocyte precursor cells (dark brown) and “4-MN” motor neurons (red). (**D**) Abundance of cluster 4-specific Elavl2 specific to the MN population (“4-MN”), GFAP for astrocyte-like cells (“1-AS”) and VAMP1 for neuronal cells (“0-N”). Secretogranin 2 (SCG2) was identified as one of the novel protein markers for in vitro generated MNs. (**E**) Expression of the corresponding mRNAs. (**F**) Summary of protein-specific GSEA of the GO:Biological Process (GO:BP) terms and Reactome pathways in MNs. (**G**) Relative organelle abundance analysis of d60 cells show differential abundance of key organelles characterized by decreased glycolysis and increased mitochondrial TCA enzymes (mt-TCA) in “4-MN”, and decreased in lysosomal and increased proteasomal abundance. (**H**) CellRank identification of terminal cell states in d52/d60 cultures. Inset represents UMAP visualization of both d52 (green) and d60 (pink) timepoints. Terminal cells states identified by CellRank: “S0” (brown), “S1-P” proliferative/neural stem cells (purple), “S2-AS” astrocytes (blue), “S3-M” cells with mixed identity (pink) and “S4-MN” motor neurons (red). (**I**) Calculated pairwise Spearman correlation coefficients for the identified terminal cell states and the SCP dataset of the developing human brain by Wu et al. (14) (OP – oligodendrocyte precursor cells, EN – excitatory neurons, uG – microglia, oRG – outer radial glia, Vasc – vasculature, IN – inhibitory neurons, I-EN – intermediate progenitor cells for EN, RG – radial glia).

After performing the uniform manifold approximation and projection (UMAP) unbedding^53^ of the proteomics data, distinct cell clusters largely corresponded to each differentiation timepoint (**Fig 2 C**). The cells’ proteomic landscape gradually moved from the pluripotent state, with downregulation of Oct4 and Lin28a (**Fig. 2 D, F**), cell cycle exit (**Fig 2 E**) and an increase in MN-enriched Nfl and glial-enriched glial fibrillary acidic protein (Gfap) at d60 (**Fig 2 D, G-H**). Additionally, a few cells deviated from the main differentiation trajectory such as rare iPSCs that clustered together with NEPI and d60 cells, as well as d60 cells that did not fully differentiate (**Fig 2 D & Sup Fig 4**).

**Figure 4.**
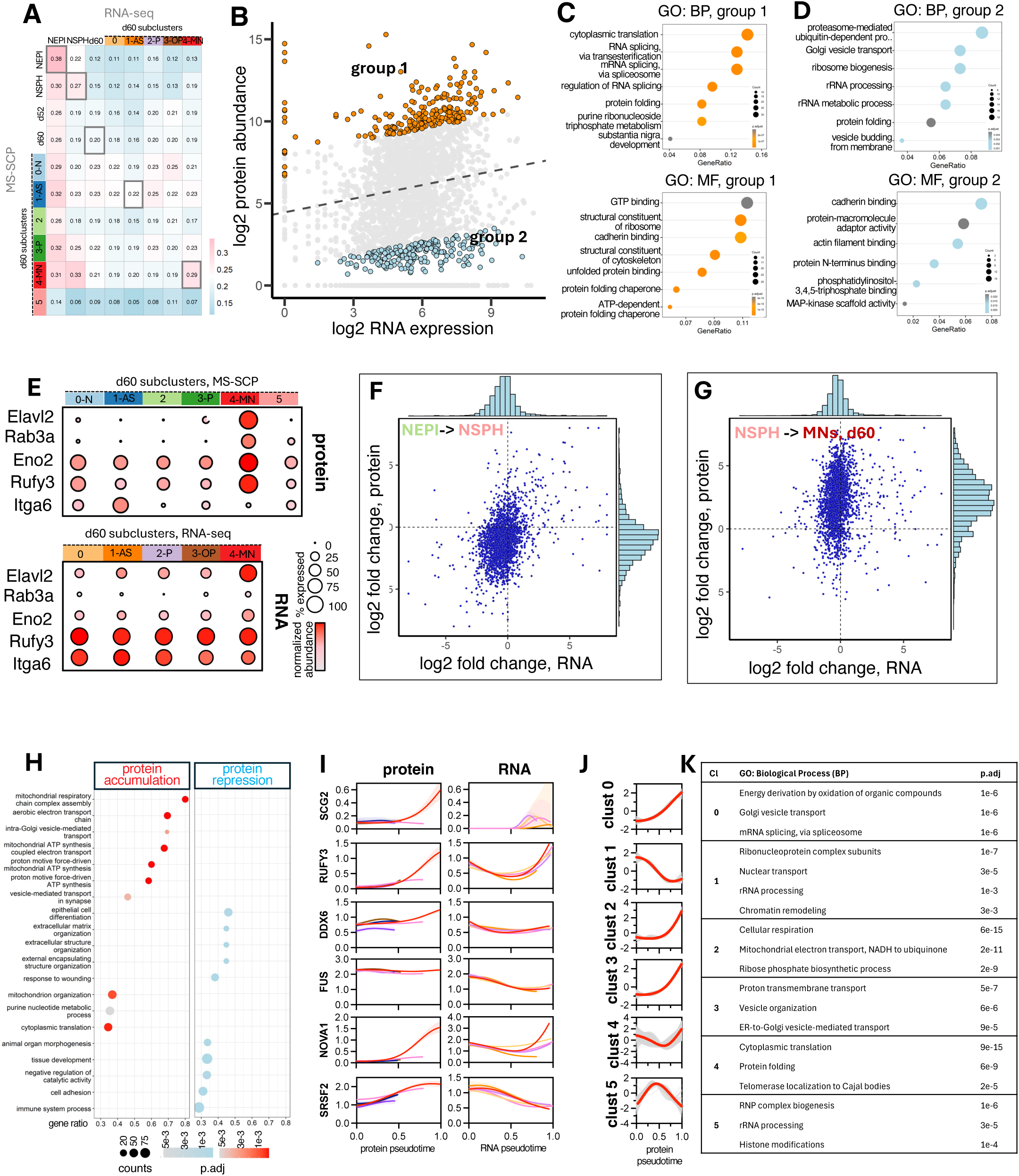
MS-SCP identifies global dynamic and cell type-specific RNA-protein discordance. (**A**) RNA expression and MS-SCP protein abundance were used to calculate pairwise Spearman correlation coefficients for MN progenitors and cellular subclusters at d60 cultures. (**B**) Protein abundance in MN cluster “4-MN” was plotted against the mRNA expression on scatter plot. X and Y axes correspond to normalized log2-transformed RNA and protein levels (log-log scale). Black line represents the linear regression line. Proteins were scored based on deviation from the regression to identify classes of proteins with high protein-low mRNA levels (group 1, orange) and low protein-high mRNA (group 2, light blue). (**C**) Identification of GO Biological Process (GO:BP, top) and Molecular Function (GO:MF, bottom) terms for high protein-low mRNA levels (group 1). (**D**) Identification of GO Biological Process (GO:BP, top) and Molecular Function (GO:MF, bottom) terms for low protein-high mRNA (group 2). (**E**) Examples of protein-RNA pairs showing MN cluster specificity and high discordance include Rab3a, Eno2, RUFY3 and Itga6 depicted on a dot plot with abundance level and % expressing cells (based on dot size). (**F**) To analyze the dynamics of RNA-protein discordance during cell differentiation from NEPI to NSPH, log2 fold change in protein level was plotted against the log2 fold change in the corresponding RNA level. (**G**) To analyze the dynamics of RNA-protein discordance during cell differentiation from NSPH to MN, log2 fold change in protein level was plotted against the log2 fold change in the corresponding RNA level. (**H**) GO:BP terms associated with proteins showing high dynamic protein accumulation during MN maturation (left, in red). GO:BP terms terms associated with protein repression during MN maturation (right, in blue). (**I**) Dynamics of protein and corresponding mRNA levels were plotted against lineage-specific pseudotime for novel MN-specific proteins (SCG2, RUFY3, DDX6) and several known RNA-binding proteins (FUS, NOVA1, SRSF2). Line colors correspond to identified CellRank states at day 60 (protein dynamics) and clusters identified by snRNA-seq (RNA dynamics).

Differentiation protein abundance analysis revealed uniquely enriched proteins at each timepoint (**Fig 2 I and Sup Table 2**). Gene set enrichment analysis (GSEA) of GO terms related to biological processes (GO:BP) showed that iPSCs had more proteins related to translation and ribonucleoprotein biogenesis, while MN progenitors (NEPI, MNP1, MNP) had enrichment in spliceosome and mRNA processing-related proteins (**Fig 2 J and Sup Table 3**)^54^. MNs at d60 had decreased ribosome processing and increased cytoskeletal organization (**Fig 2 J**). We also noticed progressive accumulation of proteins related to innate immune system (HLA-A/B/C and beta-2-microglobulin) (**Sup Fig 5 A**) that were absent in iPSCs, further confirming their immune-privileged state^55^.

Classification of proteins based on the GO cellular component (GO:CC) terms also pointed at cell type-enriched changes in organelle abundance (**Sup Table 3)**. RNA sequencing (RNA-seq) has a limited ability to approximate total mitochondrial or ribosomal abundance in cells^56^ and recent image-based approaches reveal considerable variations at the single-cell level^57^. MS-SCP now provided the opportunity to directly evaluate the structural abundance of various organelles at the single-cell level. Similar to the metabolic pathway analysis described for iPSCs, we extracted genes associated with organellar GO terms, converted them to protein names and measured normalized cumulative abundance of an organelle. Such aggregate analysis of ribosomal, endoplasmic reticulum (ER), lysosomal and mitochondrial (TCA-related) abundance revealed complex patterns of organelle re-distribution during differentiation (**Fig 2 K**). Specifically, ribosomal levels decreased as iPSCs differentiated to MNs, while the mitochondrial abundance increased in a subset of differentiated cells (**Fig 2 K and Sup Fig 5 B**). Pathway-specific scoring specifically revealed differential abundance of TCA cycle, ATP synthase and mito-ribosome proteins (**Sup Fig 5 B**). Beyond cluster-level analysis, the MS-SCP allowed us to measure organelle variations at the singe-cell level, revealing heterogeneity in abundance within clusters (**Fig 2 L and Sup Fig 5 B**). Overall, these data show complex rearrangement during MN differentiation both at the housekeeping and organelle levels as well as heterogeneity at the level of individual cells, with a potential of identifying unique cellular subtypes within a culture.

### SCP identifies various cellular types in differentiated MN cultures

To further characterize the proteomic signatures of iPSC-derived MNs we next focused on the composition of developing and more mature cultures at d52 and d60 (**Fig 3 A and B**). To aid with the cell type identification we also performed paired single-nuclei RNA sequencing (snRNA-seq) at selected timepoints on the same samples (**Fig 3 A and C**). Although we could not look at RNA and protein in the same cell, using unsupervised cluster identification algorithms, we were able to classify cells into distinct broad clusters solely based on their proteomics profile (**Fig 3 B**).

Cluster identification showed a neuronal cluster with high abundance of VAMP1 (“0-N”), astrocyte-like cells positive for GFAP (“1-AS”), proliferating neural stem cells (“3-P”) and a distinct cluster ELAVL2/3-positive Nestin (NES)-negative cells (Leiden/subpopulation “4-MN”) (**Fig 3 D**). Although canonical cholinergic neurotransmission markers such as *CHAT*, *SCL18A3* or *SLC5A7* were not detectable, likely due to their inherent low proteomic abundance and sensitivity limitations of MS-SCP, we attributed the “4-MN” cluster to putative MNs based on their marker similarity to the snRNA-seq. Specifically, snRNA-seq analysis of d60 cells similarly identified a subcluster of cells characterized by expression of *SLC5A7* and high expression of *ELAVL2/3* which are MN-enriched proteins affected in ALS (*58*) (**Fig 3 C, E and Sup Fig 6**). Consistent with the proteomics data, the transcriptomic analysis revealed an astrocyte lineage (“1-AS”) expressing GFAP/EAAT2 (**Fig 3 E and Sup Fig 6**), evidence of oligodendrocyte precursor (OP) cells expressing PDGFRA/MBP (“3-OP”) and PCNA/KI67 proliferating progenitor/neural stem cells (“2-P”).

Proteomic analysis of cells at the intermediate d52 timepoint also contained a distinct ELAVL2/3+/NES-population but lacked any glial lineages (**Sup Fig 7 A - C**), which supports immunocytochemical analysis for neuronal and glial markers in d60 cells (**Sup Fig 7 D**) and correlates with later gliogenic commitment *in vivo*^59^. Despite timepoint differences in protein abundance (**Sup Fig 7 E**), MN clusters from d52 and d60 showed very high proteomic similarity (only 17 differentially abundant proteins of ∼1654 detected) and were combined for downstream processing (**Sup Fig 7 F**). In contrast, we found that most non-MN cells at d52 and unclassified cells at d60 expressed various markers, including of neurons (NEFL, VAMP1), radial glial/astrocyte precursor cells (FABP7), and mesenchymal markers (Col1a1, Fn1) (**Sup Fig 7 G**). The transitory cells were yet to express late lineage markers that defined terminal states such as GFAP or ELAVLs (**Sup Fig 7 B, C**). These cells also expressed glioblastoma markers (CD47), and GSEA against a glioblastoma-related dataset showed enrichment in glioblastoma-related pathways for d52 cells but not the MN cluster at d60 (**Sup Fig 7 H**)^60^.

Together, this cell population at d52 and unclassified cells at d60 (**Fig 3 I**) have a fluid proteomics profile with some shared signatures of neuronal and glial cells despite expression of key neuronal transcriptional factors and neuronal-like morphological characteristics (**Fig 2 B**). Such ambiguous protein and transcriptomic profiles have also been described for less differentiated neurons states in glioblastoma^61^.

Analysis of MN-specific proteins showed enrichment of proteins related to oxidative phosphorylation, synapse maturation, folding and organelle transport (**Fig 3 F and Sup Table 4**). While some of these pathways reflected those identified by snRNA-seq (**Sup Table 5**), MS-SCP also detected unique terms related to mitochondria bioenergetics, autophagy and neuronal vesicle-mediated transport (**Fig 3 F and Sup Table 8**). We also identified proteins uniquely enriched in the MN population - specifically UCHL1 (a cortical motor neuron marker)^62^, components of cAMP-dependent protein kinase, 14-3-3 adaptor complex, as well as a number of unexpected secreted proteins and neuropeptide precursors such as VGF, secretogranin-2 (Scg2), Rufy3, and DDX6, among others (**Sup Fig 8 and Sup Tables 6 and 7**). Compared to other clusters, organelle abundance analysis revealed differential enrichment of OXPHOS proteins relative to glycolysis, and increased proteasome abundance relative to lysosomes in MNs (**Fig 3 G).**

To further refine terminal cell states in the proteomics dataset and to infer trajectory of the cells throughout their differentiation, we applied the CellRank^63^ algorithm to established temporal points (from NEPI to NSPH progenitors and d52/60 MN cultures) (**Fig 3 H and Sup Fig 9 A**). This approach identified five terminal differentiation states in d52/d60 cultures and reconstituted lineage-specific pseudotime from Palantir (**Fig 3 H and Sup Fig 9 A**). We assigned these terminal states to cell types based on the expression of canonical markers: proliferative FABP7+/PTMA+ radial glial/neural stem cells (S1-P), GFAP+/GLAST+ astrocytes (S2-AS), partially differentiated neurons described above and corresponding to day 52 cells (S3-M), as well as putative MNs (S4-MN) (**Fig 3 H and Sup Fig 9 B, C**). State S0 expressed some known markers of the OP lineage, TPM1 to 3 and Septins, which could correspond to the transcriptomic OP cluster. Cross-comparison of these clusters versus recently published SCP data from the developing human brain (*14*) showed the high degree of the proteome correlation between the *in vitro*-derived MNs and cortical excitatory neurons (EN) (R= 0.65, **Fig 3 I**) and high correlation between the astrocyte lineage and brain radial glia cells (RG) (R=0.63, **Fig 3 I**). We have also performed correlation analysis with the recently published single-cell laser-capture proteomics of human motor neurons^13^ which in contrast showed some discordance in low and mid-abundant proteins but higher concordance in high abundant proteins (**Sup Fig 10**). This discordance could be explained by the cell source (iPSC-derived versus adult) and technical platform differences.

### SCP reveals cell type-specific transcriptome-proteome discordance

How well the cellular transcriptome is predictive of the proteome is a long-standing biological question^64–66^. Such correlations between the messenger RNA (mRNA) and corresponding proteins remain variable, from moderate to strong at the whole tissue level^11,65,67^. However, less is known about their relationships at the single-cell level due to the stochastic nature of RNA transcription^68–70^ based on protein demand and post-transcriptional processing.

To look at the single-cell general correlation between individual proteins and their corresponding mRNAs at selected timepoints, we used the paired snRNA-seq and MS-SCP datasets for neuroepithelium (NEPI), neurosphere progenitors (NSPH) and differentiated states (d52/60 MNs) (**Fig 3 A**) and performed Sperman correlation analysis. We observed stronger mRNA-protein correlation for more proliferative and less differentiated NEPI cells (R=0.38) compared to NSPH (R=0.27) and day 60 cultures (R=0.20) (**Fig 4 A**). At the subcluster level, the correlations for the same pairs were highly variable for different cell types (R=0.22 for astrocytes verses R=0.29 for MNs) but higher compared to the bulk comparisons.

We then classified all mRNA-protein pairs by filtering for common features from the mRNA and protein datasets for cluster “4-MN” at d60, performing a linear regression and calculating RNA-protein Z-scores (**Fig 4 B**, log-log scale). This approach identified that outlier pairs fell into group 1 “high protein – low RNA level” (**Fig 4 B**, orange) and group 2 “low protein – high mRNA” categories (**Fig 4 B**, blue) even though, in general, this analysis revealed several magnitude-wide variabilities in individual protein levels at the same levels of mRNA. We then further classified such discordant proteins from both groups using GO. Proteins of the high protein – low mRNA group were associated with translation, splicing and folding pathways (**Fig 4 C and Supp Table 8**), while proteins of the low protein – high RNA group were associated with vesicle-mediated transport, proteasome and ribosome (**Fig 4 D**). These results correlate with the recent SCP data from the developing human cortex, where splicing pathways were the most discordant at the RNA-protein level^14^. Examples of MN-specific discordant proteins include Rab3a and neuron-specific enonalse 2 (gamma) (ENO2), while expression of ITG6A, a known astrocyte marker^71^, was high in all clusters at d60 while only detected at the protein level in cluster “1-AS” (**Fig 4 E)**. Comparing the lists for cellular subclusters showed that such discordant pairs were cell type-specific, as gene terms for MNs and astrocytes revealed both shared as well as unique pathways for each cell type (**Supp Table 8**).

To further analyze the dynamics of such unsynchronized mRNA-protein pairs, we correlated changes in RNA and protein levels between various differentiation timepoints and lineages (**Fig 4 F and G**). While cell differentiation from NEPI to NSPH revealed some discordant pairs (**Fig 4 F**), the most dramatic changes in RNA-protein correlation dynamics occurred during the progenitor neurosphere (NSPH) to MN d60 transition (**Fig 4 G**). This transition was characterized by general accumulation of numerous proteins while their mRNAs remained stable, with the density histogram showing protein amplification compared to transitions between the less differentiated states (**Fig 4 G**). Protein terms related to this dynamic accumulation mostly related to mitochondrial respiration, vesicle-mediated transport, nucleotide metabolism, and translation (**Fig 4 H, left and Sup Table 9**), consistent with the analysis of d60 MN (**Fig 3**).

For dynamic protein repression, terms were related to cell adhesion, extracellular matrix organization and integrin signaling (**Fig 4 H, right**). Individual proteins contained within these terms also included markers of proliferation (PCNA) and DNA replication (AURKB) (**Sup Table 9**). Reactivation of cell cycle genes in neurons can occur in several neurodegenerative states^72^ and our data suggest that it cannot always be predicted based solely on transcriptomics profiling. Examples of such discordant RNA and protein expression were plotted for various terminal states using MN lineage specific protein pseudotime inferred from Palantir (**Sup Fig 9 A and D**). Within each term, this analysis revealed complex regulatory patterns. Specifically, SCG2 and RUFY3 were upregulated at the protein but not at the RNA level, and RNA binding proteins DDX6, FUS, NOVA1 and SRSF2 showed discordant and concordant patterns (**Fig 4 I**). Cluster analysis of these temporal protein data revealed clusters of co-regulated MN proteins and pathways (**Fig 4 J and K and Sup Table 10)** consistent with the endpoint analysis of d60 cultures but revealing complex lineage-specific RNA-protein dynamics.

## DISCUSSION

Several groups have recently provided insights into the single-cell proteomic composition of neuronal cells *in vitro*^26,73^ and *in vivo*^13,14,25,27^. This study, to our knowledge, provides the first insights into proteomics at the single-cell level for iPSCs and iPSC-derived MNs. Many steps of post-transcriptional regulation control protein abundance and turnover^64^. Consistent with previous quantitative data^64–66^, we show considerable protein-specific discrepancies in mRNA-protein levels, but which change depending on the differentiation stage.

Importantly, our data shows that these relationships are cell type-specific, potentially making the existing protein prediction algorithms prone to error without extensive ground truth of experimental correlations in multiple cell types^74^. Unbiased determination of protein abundance allowed us to apply the MS approach to characterize a cells’ organelle abundance and metabolic states. The MS-SCP approach can now strengthen single-cell organelle measurement using antibody- and dye-based imaging approaches^57^, which has already shown considerable cell-to-cell heterogeneity of organelle abundance and can provide a future framework to relate multi-dimensional changes in organellar abundance to cell states and fates.

This methodology uncovered a surprising protein heterogeneity in iPSCs and MNs and identified protein candidates subjected to complex post-transcriptional dysregulation. Reports show that human ESCs have a higher degree of transcriptomic and surface marker homogeneity compared to human iPSCs^46^, and our data could suggest more complex post-transcriptional regulation of the pluripotent state than previously thought. Whether these regulatory mechanisms are important for its maintenance and exit and how it can impact the cell fate and terminal differentiation is a subject of future research.

Unbiased characterization of individual MNs has been recently performed across species and ALS patients^3,7,75–77^ with single-nucleus multi-omic methods (snRNA, spatial transcriptomic, and ATAC-seq). However, these RNA-only approaches miss the complexity of the MN proteomic composition and architecture and determination of genuine MN proteome and their changes remain extremely limited due to MN sparsity and signal dilution of the bulk methods^13,15^.

The ability to decipher small protein changes in rare cells at the single-cell level can be critical to understand the dynamics of cell biology and development. We predict that many proteomic changes may be missed with bulk analysis or when based solely on the transcriptome profiling. Further, changes in mRNA level can be secondary in some cases and not informative to decipher the on-mechanism of the differentiation trajectory or disease states. Finally, uncovering protein changes at the single-cell level may shed light on the role of normal protein function in neurodevelopment as well as toxic protein species accumulation and dysregulation of proteostasis in neurological conditions and injury^78^.

## ACKNOWLEDGMENTS

We thank all members of the Svendsen and Van Eyk laboratories, as well as Soshana Svendsen and Elena Vasileva for their comments and suggestions. We thank the members of the Cedars-Sinai Applied Genomics, Computation & Translational Core for their assistance in RNA sequencing and the Cedars-Sinai Proteomics and Metabolomics core for mass spectrometers access and technical support.

## AUTHOR CONTRIBUTIONS

Conceptualization: V.Z., A.B., J.V.E., C.N.S. Methodology: A.B., V.Z., A.H., J.V.E., C.N.S. Investigation: V.Z., A.B., A.H., E.I., S.B. A.S.Z., G.L. Formal analysis: V.Z., A.B. Data curation: A.B., V.Z., J.V.E., C.N.S. Visualization: V.Z., A.B., A.S.Z., G.L. Supervision: J.V.E. and C.N.S. Funding acquisition: C.N.S., J.V.E. Writing—original draft: V.Z., A.B. Writing—review and editing: V.Z., A.B., A.S.Z., J.V.E., C.N.S.

## SUPPLEMENTAL INFORMATION

Document S1. Figures S1-S10

Tables S1 to S10, Excel files containing additional data too large to fit in PDF

## METHODS

### MATERIALS AND METHODS

#### iPSC culture and motor neuron differentiation

The CS02iCTR-Tn11 human iPSC line was generated by the Cedars-Sinai Medical Center iPSC Core from peripheral blood mononuclear cells of a healthy male individual with nonintegrating oriP/EBNA1 plasmids, which allowed for episomal expression of reprogramming factors and shown to be fully pluripotent (*18*). iPSCs were maintained in mTESR1 medium on Matrigel-coated cell culture plates and passaged every 5 days at split ratios from 1:6 to 1:12 as needed using Versene. Only iPSCs between passage 17 and passage 35 were used for differentiation in this study (*18*).

The iPSCs were differentiated into motor neurons (MNs) using an established differentiation protocol utilizing small molecule morphogens (*50*). Briefly, iPSCs were cultured and expanded on Matrigel coated plates (Fisher) in mTESR medium (StemCell). Cells were dissociated with Accutase (Gibco) and plated in neuroepithelial induction medium (NEPIM: DMEM/F12:Neurobasal Plus 1:1 supplemented with B27, Glutamax, and NEAA [all from Gibco]; 0.1mM ascorbic acid [Sigma]; and 3μM CHIR99021; 2μM SB431512; and 2μM DMH1 [all from Cayman, Ann Arbor, MI, USA]) to generate neuroepithelial cells. After six days *in vitro*, neuroepithelial cells were dissociated and expanded with NEPIM containing 0.1μM retinoic acid (Sigma), 0.5μM purmorphamine (Cayman) and 0.5mM valproic acid (Sigma) to produce MN progenitors (MNPs). Next, MNPs were detached and cultured in MN induction medium (NEPIM plus 0.5μM retinoic acid, 0.1μM purmorphamine) to generate neurospheres. After six days, neurospheres were dissociated and plated on laminin-coated dishes in MN maturation medium (MN induction medium supplemented with 0.1μM Compound E [Sigma], and 20ng/ml CNTF, and 20ng/ml IGF-1, both from Pepro Tech) to induce MN differentiation. Half media changes were performed twice per week until the endpoint.

### Cell isolation and sorting

At each timepoint, cells were dissociated with Accutase, collected, and resuspended in PBS supplemented with 0.5 mM EDTA. Cells were stained for viability with Sytox Green dye (Thermo Scientific S7020, 1:5000) for 30 min on ice, washed in PBS/0.5 mM EDTA buffer, and dispensed using FACS-sorting machine (BD Biosciences), using a 100 mM nozzle and the “1.0-drop Single” sort setting with a 12/16 phase mask into separate wells of a low-binding 384-well PCR plate (Biorad HSP3801) containing 200 nl of lysis buffer (100 mM triethylammonium bicarbonate, 0.2% n-Dodecyl-β-D-maltoside, 10 ng/nl trypsin). Each experiment contained two rows of 50 cells which were used as reference for library preparation. Plates were covered with foil and stored at −80 °C for further processing.

### Sample preparation for MS-SCP

After lysis by freeze-thaw, all samples were subjected to trypsinization with 40 ng/μl trypsin, 4 h incubation at 37°C, followed by acidification with 0.1% formic acid. All seven timepoint 384-well plates were digested in the same batch before proteomic analysis using Tims-TOF single-cell proteomics. The remaining cells were pelleted and stored at -80 °C for snRNA-seq processing.

### Sample preparation for snRNA-seq

snRNA-seq of selected timepoints was performed as described elsewhere (*7*). Briefly, cell pellets of NEPI, NSPH, and d60 cultures were thawed and nuclei were isolated using gradient centrifugation on iodixanol. Nuclei integrity of DAPI-stained nuclei was verified under a confocal microscope (40× objective) before library synthesis. Good quality nuclei appear round and regular while poor quality nuclei appear shriveled or irregular shaped. No samples were excluded due to nuclei quality.

The standard 10x protocol was used per the "Chromium NextGEM Single Cell 3’ Reagent Kits v3.1 User Guide, Rev D" (single index) as described previously (*7*). The uniquely indexed libraries were pooled at equal ratio and sequenced on a NovaSeq 6000 (Illumina) as per the Single Cell 3′ v3.1 Reagent Kits User Guide, with a sequencing depth of ∼50,000 reads/cell at Cedars-Sinai Applied Genomics, Computation & Translational Core. Raw sequencing data were demultiplexed and converted to FASTQ format using bcl2fastq v2.20.

### Mass spectrometry data acquisition

The high throughput liquid chromatography set up for single-cell analysis has been previously developed by our group which involves the nanoflow dual-trap single-column configuration (nano-DTSC) (*31*). This configuration capitalized on parallelized nano-DTSC chromatography operating at 10 min of total run time per cell with peptides quantified over 8 min, thus offering an efficient solution to injection of 144 wells per day. Key modifications to the previously published article included integrating a 5 cm × 75 μm ID Ionopticks analytical column packed with 1.5 μm C18 material and interfacing it with the Bruker TimsTOF Ultra 2 spectrometer. These enhancements significantly improve throughput and sensitivity, thereby facilitating comprehensive single-cell proteomic analysis.

The analytical separation protocol entails using 0.1% formic acid in water as mobile phase A and 0.1% formic acid in acetonitrile as mobile phase B. The gradient profile begins at 9% B with a flow rate of 800 nl/min, gradually increasing to 20% B over 5 min, then to 40% B over 3.9 min. Subsequently, the flow rate ramps up to 1400 nl/min, reaching 98% B within 0.1 min, maintained for 0.5 min, followed by a drop to 9% B at 1200 nl/min over 0.05 min. The system holds at 8% B for 0.25 min before dropping the flow rate to 800 nl/min for 1 min, resulting in a total run time of 10 min.

The loading pump initiates the run by delivering mobile phase A (100% 0.1% formic acid in water) at 100 μl/min for the first 0.5 min, then gradually increasing the flow to 120 μl/min over 0.3 min, maintained for 5 min before dropping to 8 μl/min until the end of the 10-min run. The valves and trapping columns are regulated at 55°C in the Ultimate 3000 column oven compartment, while the analytical column is maintained at 60 °C using the Bruker “Toaster” oven.

In this setup, the analytical column is directly linked to a 10 μm ZDV emitter (Bruker) installed in the Bruker captive source, interfacing with the Bruker’s TimsTOF Ultra 2 mass spectrometer. The capillary voltage is set to 1900V, with dry gas flowing at 5.0 L/min and a temperature of 180 °C. Data acquisition employs DIA-PASEF, with ion accumulation and trapped ion mobility ramps set to 166 ms. DIA scans cover 90 m/z windows spanning 300 to 1200 m/z and 0.6 to 1.43 1/K0, with one full MS1 scan followed by four trapped ion mobility ramps, resulting in a cycle time of 0.86s.

### Proteomics data analysis

Data were analyzed with DIA-NN 2.3.1 Academia (Data-Independent Acquisition by Neural Networks) (*76*) using sample-specific library generated from the 50-cell pools from each iPSCs to MNs differentiation time course. Uniprot FASTA database from Dec 1st 2025 containing 20,402 protein entries (Swiss-Prot reviewed only) was used in the data processing steps. No fixed modifications and 1 variable modification (Unimod:35 - Methionine oxidation), minimum peptide length-7 amino acids, and maximum peptide length-40 amino acids; were set during the generation of the spectral library. Each search was conducted with second-pass and match-between-runs (MBR) enabled. The mass error tolerances were set at 15 ppm for both fragment and intact masses.

For analyzing the differentiation samples, the 50-cell samples DIA runs with the highest identifications were analyzed using the library-free search in DIA-NN against the complete human protein UniProt database to create a spectral library. The library-free identifications are set at <1% false discovery rate (FDR) to generate the final library by DIA-NN using the target-decoy strategy used for the analysis of all time points.

### MS-SCP data analysis

Analysis of MS-SCP was performed as previously described (*18*) in Seurat with the following modifications for proteomics. Cells underwent quality filtering based on total intensity (> 100,000 total abundance per cell) and unique protein count (> 150/cell). Wells containing large quantities (> 6%) of junk proteins (such as cellular keratins or epidermal proteins) were also filtered out prior to further analysis. After that, missing proteins were imputed at 0.1x minimal detected value and their abundance was normalized by total abundance per well. Values were log2(x+1) transformed. Highly variable proteins were selected based on variance-stabilized transformation in Seurat:: FindVariableFeatures. All features were used for principal component (PC) analysis to calculate a neighborhood graph with n=20 components. Calculated PCs were used for subsequent UMAP embedding in Seurat. Unsupervised cell clustering was performed using the Leiden algorithm. Clusters were manually annotated using the abundance of marker genes.

Differential expressed proteins in each cluster were calculated by comparing protein abundance in each cluster against all other clusters by Seurat:: FindAllMarkers() command with Wilcoxon rank-sum test. P-values were corrected via the Benjamini–Hochberg procedure and a cutoff of 5% FDR was applied. For Gene Onthology (GO) analysis, markers were filtered by adjusted p_value < 0.05 and only positive markers were considered. For gene set enrichment analysis (GSEA) markers were ranked based on their log fold changed. GSEA was performed on sorted log2 fold change rank lists for all proteins identified in each cluster. GO and GSEA analyses were performed in ClusterProfiler and terms were simplified before plotting. GSEA gene sets for GO biological process (BP), cellular component (CC) and molecular function (MF) with minimum set size of 3 and maximum of 1000 were used for computation and visualization. Volcano plots were generated in GraphPad Prism.

Organelle scored were calculated by first selecting proteins corresponding to the following structural GO terms: GO:0005838 (19S proteasome), GO:0005789 (ER membrane proteins), GO:0044391 (ribosomal proteins), GO:0006096 (glycolysis proteins), GO:0061621 (lysosomal proteins). Mitochondrial proteins were selected from MitoCarta3.0. To calculate the combined organellar or pathway scores for each cell, the combined score was calculated by dividing the sum of individual proteins in each GO term by total protein abundance. Timepoint specific metrics were calculated by dividing average term abundance by average total abundance in cluster. For mitochondrial pathways, pathway-specific abundances were calculated for each pathway in MitoCarta3.0 (*80*). Benjamini-Hochberg (BH) correction was used to calculate adjusted p-values.

Cell cycle scoring was performed essentially as described before using CellCycleScoring() (*78*). When needed, gene symbols, Uniprot IDs and ENTREZ gene IDs were interconverted in human AnnotationDbi (*79*). Comparison with Ai et al study (*18*) was performed by selecting common detected proteins and integrating in Harmony (*33*) using standard parameters.

CellRank analysis (*61*) was performed as described with standard setting applied. NEPI cells were defined as a starting (root) population and pseudotime was calculated using Temporal Kernel. Subsequently, a transition matrix was computed and macrostates were determined via GPCCA. Terminal states were determined by fitting six macrostates to a Schur decomposition and selecting the top five. Subsequently, MN subcluster 4 was manually added as additional terminal states. Cluster Protein Palantir pseudotime was calculated by taking terminal states as end populations and NEPI cells as root (*80*). For RNA Palantir pseudotime endpoints were defined as Seurat subclusters and NEPI cells as a root. markers were identified by Wilcoxon rank-sum test.

### snRNA-seq data analysis

Demultiplexed fastq files were run via CellRanger v6.1.2 using the "cellranger count" command with the "--include-introns" option using the precompiled CellRanger human reference sequence “v2020-A” (10x Genomics) (*81*). Expression matrices were combined into a single object using Seurat in R (*82*) and filtered by removing all cells with counts, genes, mitochondrial gene percentage, and ribosomal gene percentage outside of three standard deviations from the mean. The data was normalized, single-cell transformed and scaled. Principal component analysis (PCA) was run on the object using 14 principal components (PCs). The number of PCs was identified using an elbow plot analysis. Next, the nuclei were clustered using the Seurat functions “RunUMAP()”, “FindNeighbors()”, and “FindClusters()”. A resolution factor was adjusted to provide reasonably defined clusters for each of 0.3 identified 5 distinct clusters. Known cell type markers were used to assign general cell types to clusters. Astrocytes (AS): GFAP, EAAT1; OP cells: PDGFRA, MBP; motor neurons (MNs): CHAT, SLC5A7, ELAVL2/3, neurons: NFL and VAMP1.

Gene set enrichment analysis was conducted on differentially expressed genes and simplified using ClusterProfiler (*83*) package in R with GO biological process, molecular function, and cellular component gene sets.

### Correlation analysis between SCP-MS and snRNA-seq

RNA-protein correlation analysis was performed by first calculating pseudobulk RNA expression or protein abundance in Seurat-identified clusters and samples. These values were normalized by library size and total protein abundance per sample and log2 transformed. Z-scores were derived by calculating protein-RNA difference and scaling. Discordant RNA-protein pairs were selected by having Z-scores > 1 (group 1) or < -1 (group 2). Pathway analysis was performed on these proteins as described above.

To calculate changes and concordance in RNA-protein levels during cell differentiation, we calculated pairwise log2 fold changes in RNA and proteins levels between neurospheres and neuroepithelial cells, and between putative MNs and neurospheres. Scatter plots and density histograms were constructed in R with ggMarginal() command.

### Immunocytochemistry

For immunocytochemistry of Day-0 iPSCs, Cy3-conjugated TRA-1-60 primary antibody (MAB4360C3, Sigma-Aldrich) was diluted 1:100 with mTeSR Plus culture medium and added to live cells for 30 minutes at 37°C and 5% CO_2_. Cells were washed two times with mTeSR Plus media. Cells were imaged using a Keyence BZ-X810 fluorescence microscope.

For immunocytochemistry of d52 and d60 cultures, cells on glass coverslips were fixed with 4% paraformaldehyde (Sigma) for 30 min, permeabilized with 0.5% Triton X-100 and incubated for 30 min with blocking solution (0.5% Triton X-100,10% BSA in PBS). Cells were incubated with primary antibodies against MNX1/Hb9 (1:300) (Abcam, #ab221884), Nfh (1:1000) (Millipore-Sigma, #AB5539), ChAT (1:300) (R&D Systems, # AF3447), and GFAP (1:1000) (Agilent Dako, #Z033401-2) followed by species-appropriate Alexa Fluor–conjugated secondary antibodies. Nuclei were counterstained with 4′,6-diamidino-2-phenylindole (DAPI; Thermo Fisher Scientific).

### Immunocytochemistry quantification

Cell population analyses were generated from confocal images acquired on a Nikon A1R microscope using a 60× oil immersion objective. For each condition, multiple non-overlapping fields were imaged per coverslip. Cell quantification was performed using Fiji (ImageJ) with a custom macro to enable automated and unbiased image analysis.

Nuclei were identified based on DAPI staining using automated thresholding and watershed segmentation to separate closely apposed nuclei. Total cell number was defined as the number of DAPI-positive nuclei per field. Marker-positive cells (Nfh, MNX1/Hb9 and ChAT) were identified by applying intensity thresholds to the corresponding fluorescence channels and scoring nuclei that overlapped with marker-positive signal. For each image, marker-positive signal was identified using intensity thresholding, and the proportion of the image field occupied by GFAP-positive signal was calculated relative to the total image area. Threshold parameters were kept constant across all conditions within an experiment. Automated segmentation was visually inspected for accuracy, and parameters were kept constant across all conditions.

### xxFigure design and visualization

Graphical illustrations depicting study overview and cell differentiation were designed and prepared with Biorender.com and Adobe Illustrator. CellRank analysis was performed in Scanpy. Graphs, heatmaps, and other plots were generated with R and GraphPad Prism and assembled in Adobe Illustrator.

## REFERENCES

1. S. Sances et al., Modeling ALS with motor neurons derived from human induced pluripotent stem cells. Nat Neurosci 19, 542–553 (2016).

2. S. P. Pasca et al., A framework for neural organoids, assembloids and transplantation studies. Nature 639, 315–320 (2025).

3. O. Gautier et al., Challenges of profiling motor neuron transcriptomes from human spinal cord. Neuron 111, 3739–3741 (2023).

4. A. Yadav et al., A cellular taxonomy of the adult human spinal cord. Neuron 111, 328–344 e327 (2023).

5. R. Ho et al., Cross-Comparison of Human iPSC Motor Neuron Models of Familial and Sporadic ALS Reveals Early and Convergent Transcriptomic Disease Signatures. Cell Syst 12, 159–175 e159 (2021).

6. D. Zhang et al., Spatial transcriptomics and single-nucleus RNA sequencing reveal a transcriptomic atlas of adult human spinal cord. Elife 12, (2024).

7. J. Andersen et al., Single-cell transcriptomic landscape of the developing human spinal cord. Nat Neurosci 26, 902–914 (2023).

8. T. Rayon, R. J. Maizels, C. Barrington, J. Briscoe, Single-cell transcriptome profiling of the human developing spinal cord reveals a conserved genetic programme with human-specific features. Development 148, (2021).

9. E. G. Baxi et al., Answer ALS, a large-scale resource for sporadic and familial ALS combining clinical and multi-omics data from induced pluripotent cell lines. Nat Neurosci 25, 226–237 (2022).

10. L. C. Neuro et al., An integrated multi-omic analysis of iPSC-derived motor neurons from C9ORF72 ALS patients. iScience 24, 103221 (2021).

11. B. C. Carlyle et al., A multiregional proteomic survey of the postnatal human brain. Nat Neurosci 20, 1787–1795 (2017).

12. W. Xu et al., Machine learning-based proteomics profiling of ALS identifies downregulation of RPS29 that maintains protein homeostasis and STMN2 level. Commun Biol 8, 1177 (2025).

13. A. J. Guise et al., TDP-43-stratified single-cell proteomics of postmortem human spinal motor neurons reveals protein dynamics in amyotrophic lateral sclerosis. Cell Rep 43, 113636 (2024).

14. T. Wu et al., Single-cell proteomic landscape of the developing human brain. Nat Biotechnol, (2026).

15. Y. Cong et al., Ultrasensitive single-cell proteomics workflow identifies >1000 protein groups per mammalian cell. Chem Sci 12, 1001–1006 (2020).

16. S. Kreimer et al., High-Throughput Single-Cell Proteomic Analysis of Organ-Derived Heterogeneous Cell Populations by Nanoflow Dual-Trap Single-Column Liquid Chromatography. Anal Chem 95, 9145–9150 (2023).

17. L. Ai et al., Single-Cell Proteomics Reveals Specific Cellular Subtypes in Cardiomyocytes Derived From Human iPSCs and Adult Hearts. Mol Cell Proteomics 24, 100910 (2025).

18. A. A. Petelski et al., Multiplexed single-cell proteomics using SCoPE2. Nat Protoc 16, 5398–5425 (2021).

19. B. Furtwangler et al., Mapping early human blood cell differentiation using single-cell proteomics and transcriptomics. Science 390, eadr8785 (2025).

20. J. Derks et al., Increasing the throughput of sensitive proteomics by plexDIA. Nat Biotechnol 41, 50–59 (2023).

21. P. Su et al., Highly multiplexed, label-free proteoform imaging of tissues by individual ion mass spectrometry. Sci Adv 8, eabp9929 (2022).

22. A. Leduc, L. Khoury, J. Cantlon, S. Khan, N. Slavov, Massively parallel sample preparation for multiplexed single-cell proteomics using nPOP. Nat Protoc 19, 3750–3776 (2024).

23. H. Zhang et al., TEMI: tissue-expansion mass-spectrometry imaging. Nat Methods 22, 1051–1058 (2025).

24. S. S. Rubakhin, E. V. Romanova, P. Nemes, J. V. Sweedler, Profiling metabolites and peptides in single cells. Nat Methods 8, S20–29 (2011).

25. C. C. Johnson et al., Proteome-Driven Phenotyping of Identified Single Neurons in Intact Brain Tissue by Aspiration Patch Proteomics. bioRxiv, (2026).

26. S. Ghatak et al., Single-Cell Patch-Clamp/Proteomics of Human Alzheimer’s Disease iPSC-Derived Excitatory Neurons Versus Isogenic Wild-Type Controls Suggests Novel Causation and Therapeutic Targets. Adv Sci (Weinh) 11, e2400545 (2024).

27. L. Rodriguez et al., Patch-Clamp Single-Cell Proteomics in Acute Brain Slices: A Framework for Recording, Retrieval, and Interpretation. ACS Chem Neurosci 17, 1303–1315 (2026).

28. B. Chazarin et al., SCarP: Proteome Heterogeneity Characterization of Primary Mouse Cardiomyocytes. Circ Res 137, 1377–1379 (2025).

29. M. J. Workman et al., Large-scale differentiation of iPSC-derived motor neurons from ALS and control subjects. Neuron 111, 1191–1204 e1195 (2023).

30. V. Volpato et al., Reproducibility of Molecular Phenotypes after Long-Term Differentiation to Human iPSC-Derived Neurons: A Multi-Site Omics Study. Stem Cell Reports 11, 897–911 (2018).

31. C. R. Bye et al., Large-scale drug screening in iPSC-derived motor neurons from sporadic ALS patients identifies a potential combinatorial therapy. Nat Neurosci 29, 40–52 (2026).

32. P. Cahan, G. Q. Daley, Origins and implications of pluripotent stem cell variability and heterogeneity. Nat Rev Mol Cell Biol 14, 357–368 (2013).

33. S. Kreimer et al., Parallelization with Dual-Trap Single-Column Configuration Maximizes Throughput of Proteomic Analysis. Anal Chem 94, 12452–12460 (2022).

34. Y. Wang et al., Liquid Chromatographic and Mass Spectrometric Methods for Quantitative Proteomic Analysis from Single-Cell and Nanogram-Level Samples. Anal Chem 97, 18415–18422 (2025).

35. I. Korsunsky et al., Fast, sensitive and accurate integration of single-cell data with Harmony. Nat Methods 16, 1289–1296 (2019).

36. M. F. Pera, B. Reubinoff, A. Trounson, Human embryonic stem cells. J Cell Sci 113 ( Pt 1), 5–10 (2000).

37. A. Leduc, R. G. Huffman, J. Cantlon, S. Khan, N. Slavov, Exploring functional protein covariation across single cells using nPOP. Genome Biol 23, 261 (2022).

38. B. T. Jackson, L. W. S. Finley, Metabolic regulation of the hallmarks of stem cell biology. Cell Stem Cell 31, 161–180 (2024).

39. R. Aviner, A. Shenoy, O. Elroy-Stein, T. Geiger, Uncovering Hidden Layers of Cell Cycle Regulation through Integrative Multi-omic Analysis. PLoS Genet 11, e1005554 (2015).

40. C. Rega et al., High resolution profiling of cell cycle-dependent protein and phosphorylation abundance changes in non-transformed cells. Nat Commun 16, 2579 (2025).

41. L. Vallier, Cell Cycle Rules Pluripotency. Cell Stem Cell 17, 131–132 (2015).

42. K. A. Gonzales et al., Deterministic Restriction on Pluripotent State Dissolution by Cell-Cycle Pathways. Cell 162, 564–579 (2015).

43. S. Pauklin, L. Vallier, The cell-cycle state of stem cells determines cell fate propensity. Cell 155, 135–147 (2013).

44. H. Niwa, J. Miyazaki, A. G. Smith, Quantitative expression of Oct-3/4 defines differentiation, dedifferentiation or self-renewal of ES cells. Nat Genet 24, 372–376 (2000).

45. D. Strebinger et al., Endogenous fluctuations of OCT4 and SOX2 bias pluripotent cell fate decisions. Mol Syst Biol 15, e9002 (2019).

46. K. H. Narsinh et al., Single cell transcriptional profiling reveals heterogeneity of human induced pluripotent stem cells. J Clin Invest 121, 1217–1221 (2011).

47. A. A. Kolodziejczyk et al., Single Cell RNA-Sequencing of Pluripotent States Unlocks Modular Transcriptional Variation. Cell Stem Cell 17, 471–485 (2015).

48. T. Kalmar et al., Regulated fluctuations in nanog expression mediate cell fate decisions in embryonic stem cells. PLoS Biol 7, e1000149 (2009).

49. C. Y. Fong, G. S. Peh, K. Gauthaman, A. Bongso, Separation of SSEA-4 and TRA-1-60 labelled undifferentiated human embryonic stem cells from a heterogeneous cell population using magnetic-activated cell sorting (MACS) and fluorescence-activated cell sorting (FACS). Stem Cell Rev Rep 5, 72–80 (2009).

50. A. Filipczyk et al., Network plasticity of pluripotency transcription factors in embryonic stem cells. Nat Cell Biol 17, 1235–1246 (2015).

51. J. P. Saxe, A. Tomilin, H. R. Scholer, K. Plath, J. Huang, Post-translational regulation of Oct4 transcriptional activity. PLoS One 4, e4467 (2009).

52. Z. W. Du et al., Generation and expansion of highly pure motor neuron progenitors from human pluripotent stem cells. Nat Commun 6, 6626 (2015).

53. E. Becht et al., Dimensionality reduction for visualizing single-cell data using UMAP. Nat Biotechnol, (2018).

54. Y. Hao et al., Temporal proteomic and phosphoproteomic dynamics during neuronal differentiation in the reference iPSC line KOLF2.1J. Sci Signal 19, eadx8680 (2026).

55. K. Koga, B. Wang, S. Kaneko, Current status and future perspectives of HLA-edited induced pluripotent stem cells. Inflamm Regen 40, 23 (2020).

56. J. P. Rooney et al., PCR based determination of mitochondrial DNA copy number in multiple species. Methods Mol Biol 1241, 23–38 (2015).

57. S. Seal et al., Cell Painting: a decade of discovery and innovation in cellular imaging. Nat Methods 22, 254–268 (2025).

58. S. Diaz-Garcia et al., Nuclear depletion of RNA-binding protein ELAVL3 (HuC) in sporadic and familial amyotrophic lateral sclerosis. Acta Neuropathol 142, 985–1001 (2021).

59. D. H. Rowitch, A. R. Kriegstein, Developmental genetics of vertebrate glial-cell specification. Nature 468, 214–222 (2010).

60. R. G. Verhaak et al., Integrated genomic analysis identifies clinically relevant subtypes of glioblastoma characterized by abnormalities in PDGFRA, IDH1, EGFR, and NF1. Cancer Cell 17, 98–110 (2010).

61. J. J. Loh, S. Ma, Hallmarks of cancer stemness. Cell Stem Cell 31, 617–639 (2024).

62. B. Genc et al., Upper motor neurons are a target for gene therapy and UCHL1 is necessary and sufficient to improve cellular integrity of diseased upper motor neurons. Gene Ther 29, 178–192 (2022).

63. P. Weiler, M. Lange, M. Klein, D. Pe’er, F. Theis, CellRank 2: unified fate mapping in multiview single-cell data. Nat Methods 21, 1196–1205 (2024).

64. Y. Liu, A. Beyer, R. Aebersold, On the Dependency of Cellular Protein Levels on mRNA Abundance. Cell 165, 535–550 (2016).

65. B. Schwanhausser et al., Global quantification of mammalian gene expression control. Nature 473, 337–342 (2011).

66. C. Vogel, E. M. Marcotte, Insights into the regulation of protein abundance from proteomic and transcriptomic analyses. Nat Rev Genet 13, 227–232 (2012).

67. M. Murgas, P. Michenthaler, M. B. Reed, G. Gryglewski, R. Lanzenberger, Correlation of receptor density and mRNA expression patterns in the human cerebral cortex. Neuroimage 256, 119214 (2022).

68. Y. Taniguchi et al., Quantifying E. coli proteome and transcriptome with single-molecule sensitivity in single cells. Science 329, 533–538 (2010).

69. Y. R. Jiang et al., Simultaneous deep transcriptome and proteome profiling in a single mouse oocyte. Cell Rep 42, 113455 (2023).

70. J. M. Fulcher et al., Parallel measurement of transcriptomes and proteomes from same single cells using nanodroplet splitting. Nat Commun 15, 10614 (2024).

71. L. Barbar et al., CD49f Is a Novel Marker of Functional and Reactive Human iPSC-Derived Astrocytes. Neuron 107, 436–453 e412 (2020).

72. D. Wu, J. K. Sun, K. H. Chow, Neuronal cell cycle reentry events in the aging brain are more prevalent in neurodegeneration and lead to cellular senescence. PLoS Biol 22, e3002559 (2024).

73. B. Shen et al., Capillary Electrophoresis Mass Spectrometry for Scalable Single-Cell Proteomics. Front Chem 10, 863979 (2022).

74. M. Inayatullah, A. K. Dwivedi, V. K. Tiwari, Advances in single-cell omics: Transformative applications in basic and clinical research. Curr Opin Cell Biol 95, 102548 (2025).

75. J. A. Blum et al., Single-cell transcriptomic analysis of the adult mouse spinal cord reveals molecular diversity of autonomic and skeletal motor neurons. Nat Neurosci 24, 572–583 (2021).

76. L. M. Gittings et al., Cryptic exon detection and transcriptomic changes revealed in single-nuclei RNA sequencing of C9ORF72 patients spanning the ALS-FTD spectrum. Acta Neuropathol 146, 433–450 (2023).

77. J. Humphrey et al., Integrative transcriptomic analysis of the amyotrophic lateral sclerosis spinal cord implicates glial activation and suggests new risk genes. Nat Neurosci 26, 150–162 (2023).

78. A. Hohn, A. Tramutola, R. Cascella, Proteostasis Failure in Neurodegenerative Diseases: Focus on Oxidative Stress. Oxid Med Cell Longev 2020, 5497046 (2020).

79. V. Demichev et al., dia-PASEF data analysis using FragPipe and DIA-NN for deep proteomics of low sample amounts. Nat Commun 13, 3944 (2022).

80. S. Rath et al., MitoCarta3.0: an updated mitochondrial proteome now with sub-organelle localization and pathway annotations. Nucleic Acids Res 49, D1541–D1547 (2021).

81. I. Tirosh et al., Dissecting the multicellular ecosystem of metastatic melanoma by single-cell RNA-seq. Science 352, 189–196 (2016).

82. C. M. Pagès H., Falcon S., Li N., AnnotationDbi: Manipulation of SQLite-based annotations in Bioconductor. doi:10.18129/B9.bioc.AnnotationDbi R package version 1.74.0, https://bioconductor.org/packages/AnnotationDbi., (2026).

83. M. Setty et al., Characterization of cell fate probabilities in single-cell data with Palantir. Nat Biotechnol 37, 451–460 (2019).

84. G. X. Zheng et al., Massively parallel digital transcriptional profiling of single cells. Nat Commun 8, 14049 (2017).

85. Y. Hao et al., Integrated analysis of multimodal single-cell data. Cell 184, 3573–3587 e3529 (2021).

86. G. Yu, L. G. Wang, Y. Han, Q. Y. He, clusterProfiler: an R package for comparing biological themes among gene clusters. OMICS 16, 284–287 (2012).

